# Oleic acid decouples fecundity and longevity via DAF-12 steroid hormone signaling in *C. elegans*

**DOI:** 10.1101/2023.04.13.536759

**Authors:** Alexandra M. Nichitean, Frances V. Compere, Sarah E. Hall

**Affiliations:** Department of Biology, Syracuse University, Syracuse, NY 13244

**Keywords:** *C. elegans*, postdauer, DAF-12, FAT-7, VIT-2, starvation, dauer, oleic acid, ferroptosis

## Abstract

In animals, early-life starvation can program gene expression changes that result in profound effects on adult phenotypes. For *C. elegans* nematodes, passage through the stress-resistant dauer diapause stage due to early-life starvation establishes a cellular memory that manifests as increased metabolism and decreased fecundity compared to continuously developed adults. To further investigate the connection between metabolism and reproduction, we supplemented the diet of postdauer adults with different fatty acids and examined their life history traits. Here, we show that dietary oleic acid (OA) supplementation uniquely increases the fecundity of both postdauer and continuously developed adults in a DAF-12 steroid signaling dependent manner, potentially through the increased expression of *fat-7* Δ9-desaturase and *vit-2* vitellogen genes. In addition, OA may rescue increased ferroptosis in postdauer germ lines and has complex effects on adult lifespan depending on the animals’ life history. Together, our results suggest a model where OA modifies DAF-12 activity to positively regulate fecundity, metabolism, and lifespan in adults.

## Introduction

Undernutrition *in utero* or early life is associated with increased risk of adult disease in humans. For example, infants born during the Dutch Hunger Winter (1944-45) exhibited low birth weights and had higher incidences of obesity, lower glucose tolerances, and cardiovascular disease in adulthood compared to siblings not born during the famine (L. H. Lumey and Stein 1997; Portrait, Teeuwiszen, and Deeg 2011; Kahn, Stein, and Lumey 2010; L. Lumey et al. 2009; Kahn et al. 2009; Yarde et al. 2013). These adverse health outcomes have persisted for multiple generations despite the absence of direct nutritional stress to those individuals (“Transgenerational Effects of Prenatal Exposure to the 1944–45 Dutch Famine - Veenendaal - 2013 - BJOG: An International Journal of Obstetrics & Gynaecology - Wiley Online Library” n.d.). Additionally, the Chinese famine (1959-61) resulted in similar health outcomes, with severe starvation stress *in utero* correlating with increased risk of developing symptoms associated with type 2 diabetes, such as hyperglycemia (Li et al. 2010). Together, these observations indicate that nutritional stress during critical periods of development triggers a cellular ”starvation memory” that persists into adulthood to alter metabolic phenotypes.

To investigate the molecular nature of this cellular starvation memory, we utilized the natural developmental decision made by *Caenorhabditis elegans* nematodes based on their environmental conditions in early life. When cultivated in favorable growth conditions, *C. elegans* animals proceed continuously through four larval molts before becoming reproductive adults (controls or CON adults) (Sulston and Horvitz 1977). Alternatively, L1 larval stage animals exposed to environmental stress, such as starvation, during a critical period are driven to enter the alternate dauer diapause stage. Dauer larvae are developmentally quiescent, stress resistant, non-feeding, and non-aging. If conditions improve or they relocate to a more favorable environment, dauer larvae will resume continuous development to become reproductive postdauer (PD_Stv_) adults (Cassada and Russell 1975). We showed previously that postdauer adults retain a cellular memory of their developmental history that is specific to the dauer inducing stress (Ow et al. 2018). For postdauer adults that experienced early-life starvation, somatically-expressed metabolic genes and germline-expressed reproductive genes were up and down-regulated, respectively, compared to control adults. These changes in gene expression correlated with altered life history traits. For example, PD_Stv_ adults stored less lipids in their intestines, likely due to the increased mobilization of lipids from the intestine into oocytes, a process known as vitellogenesis (Ow, Nichitean, and Hall 2021; Kimble and Sharrock 1983; Grant and Hirsh 1999). These animals also exhibited significant decreases in progeny number, partially resulting from a delay in gametogenesis after dauer rescue, which correlated with an increase in longevity compared to controls (Ow, Nichitean, and Hall 2021). Interestingly, F1 progeny of PD_Stv_ adults inherited the “starvation memory”, resulting in the increased storage of intestinal lipids compared to the F1 progeny of control adults. Our results demonstrated that the response to starvation stress experienced during critical periods of development is conserved in animals, resulting in altered metabolic phenotypes that can be inherited similar to humans.

Furthermore, the altered life history traits we characterized in PD_Stv_ adults requires the somatic aging pathways, including the DAF-12 steroid signaling pathway (Ow, Nichitean, and Hall 2021). DAF-12 is a nuclear hormone receptor with homology to the vitamin D receptor and is known as the “master regulator” due to its role in regulating aging, metabolism, developmental timing, and dauer formation (Antebi et al. 2000; Hochbaum et al. 2011). When bound to its ligand, DAF-12 promotes expression of genes that have functions in reproductive development. The ligands, dafachronic acids (DA), are synthesized from dietary cholesterol in part by DAF-36 Rieske oxygenase and the HSD-1 and DHS-16 dehydrogenases (Rottiers et al. 2006; Mahanti et al. 2014; Patel et al. 2008; Wollam et al. 2012). DAF-9, a cytochrome p450, regulates the last step in the DA synthesis (Motola et al. 2006). Multiple types of DA have been isolated and have been shown to differentially modulate DAF-12 activity when bound as a ligand (Mahanti et al. 2014). We have shown that DA-dependent DAF-12 activity regulates the onset of germline proliferation, progeny number, lipid storage, and vitellogenesis in PDStv adults, suggesting that passage through dauer alters DAF-12 activity and/or gene targets to alter life history traits based on developmental experience (Ow, Nichitean, and Hall 2021).

To further investigate the connection between the altered metabolism and reproduction due to starvation-induced dauer formation, we previously tested the effects of supplementing the diet of PD_Stv_ animals with the mono-unsaturated fatty acid, oleic acid (OA). Unexpectedly, we observed that OA increased the progeny number of PD_Stv_ animals, rescuing the decreased fecundity on a normal diet. This OA-dependent increase in fecundity required the Δ9 desaturase FAT-7 that functions in the fatty acid metabolism pathway, but not NHR-49/HNF4 transcription factor that regulates the expression of desaturase genes (Ow, Nichitean, and Hall 2021). Here, we further explore the mechanism of the OA-dependent increase in fecundity by showing that CON adults also show this phenotype on an OA-supplemented diet, and that DAF-12, DAF-9, and HSD-1 are required in both populations for the effect. We show that OA supplementation promotes increased expression of *fat-7* and the vitellogen gene, *vit-2*, in a DAF-12 dependent manner. We provide evidence that OA acts late in development to alter progeny number, and it may be acting to rescue an increased level of ferroptosis specifically in PD_Stv_ to increase brood size. Finally, we demonstrate the OA increases longevity of both CON and PD_Stv_ adults, showing that the trade-off between reproduction and lifespan can be de-coupled through OA supplementation. Together, our results indicate that OA improves multiple life history traits in *C. elegans* adults by altering DAF-12 activity and rescuing physiological ferroptosis in the germ line.

## RESULTS

### Oleic acid uniquely increases fecundity in *C. elegans* adults

We showed previously that dietary supplementation with oleic acid (OA) throughout life rescued the decreased brood size phenotype observed in PD_Stv_ adults (Ow, Nichitean, and Hall 2021). To investigate whether oleic acid was alleviating nutrition deficits due to starvation in early development, we measured the brood size of PD_Stv_ animals whose diet was supplemented with a variety of mono and poly-unsaturated fatty acids (MUFAs and PUFAs, respectively) representing different stages of fatty acid metabolism, including palmitoleic acid, linoleic acid, arachidonic acid, and eicopentaenoic acid (Figure 1A). In *C. elegans*, acetyl-CoA is converted into the saturated fatty acid, palmitic acid, which be modified in two different pathways. In one path, palmitic acid can be modified by the FAT-5 desaturase to form the MUFA palmitoleic acid (Watts and Browse 2000). Alternatively, palmitic acid can be converted by elongase enzymes into stearic acid, after which various desaturases, including FAT-6, FAT-7, and FAT-2, can catalyze the formation of MUFA OA and many types of PUFAs (Figure 1A) (reviewed in (Watts 2016)). As we reported previously, dietary supplementation with OA throughout the life of PD_Stv_ adults resulted in significantly increased brood sizes compared to PD_Stv_ adults fed a control diet. However, dietary supplementation with palmitoleic acid, linoleic acid, arachidonic acid, and eicopentaenoic acid had no significant effects on the brood size of PD_Stv_ adults (Figure 1B). These results suggest that lower amounts of available fatty acids in PD_Stv_ adults, per se, are not responsible for their decreased brood size, and that OA is special in its ability to affect their reproductive capacity.

**Figure 1.**
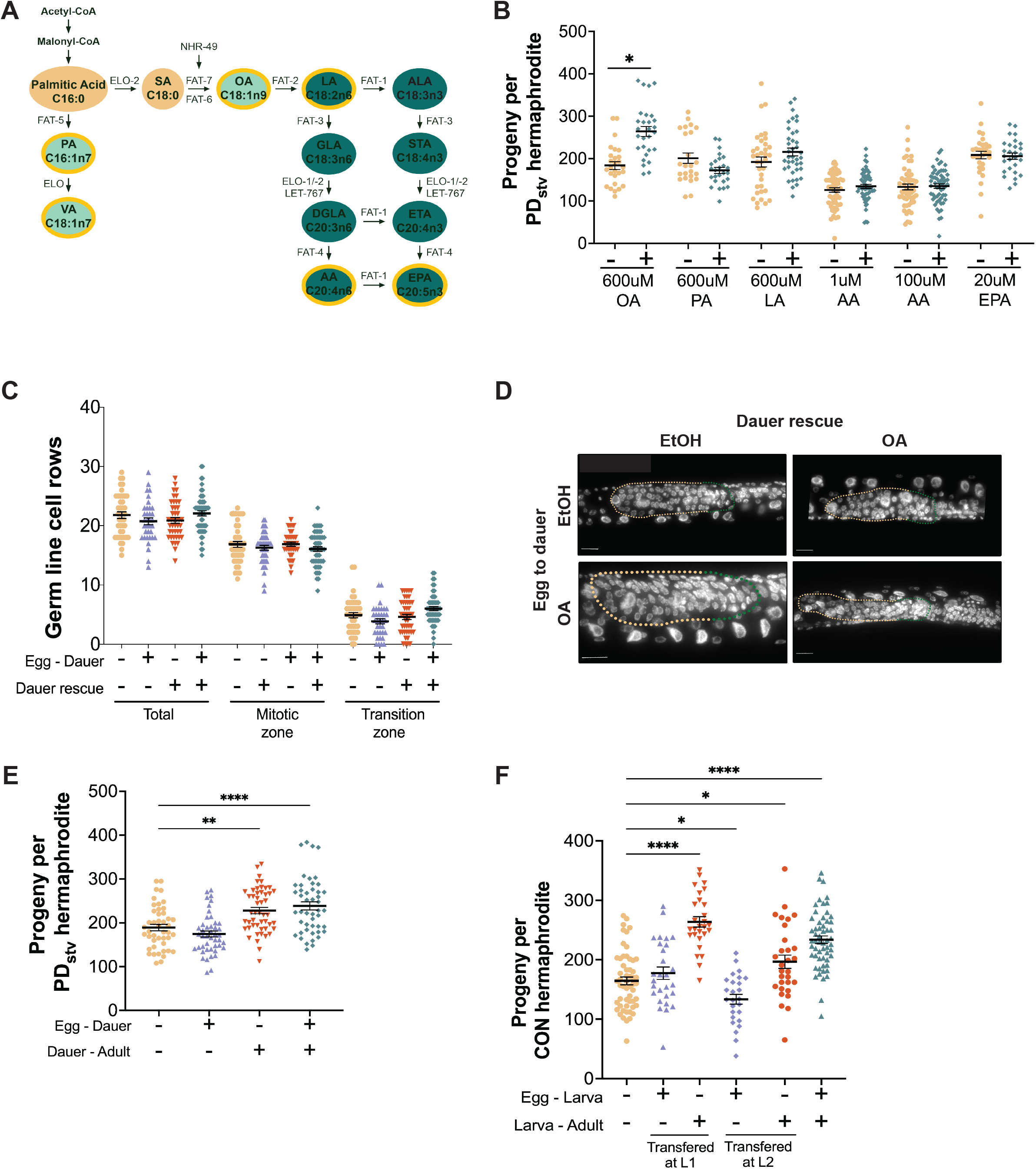
Oleic acid increases fecundity in *C. elegans* adults. (**A**) Model of fat metabolism pathway in *C. elegans*. Saturated fatty acids (tan): SA, stearic acid. Monounsaturated fatty acids (light green): OA, oleic acid; PA, palmitoleic acid; VA, vaccenic acid. Polyunsaturated fatty acids (dark green): LA, linoleic acid; ALA, α-linoleic acid; GLA, γ-linoleic acid; STA, steatidonic acid; DGLA, dihomo-γ-linoleic acid; ETA, eicosatrienoic acid; AA, arachidonic acid; EPA, eicosapentaenoic acid. Fatty acids used in (**B**) are circled in yellow. (**B**) Brood sizes of wildtype PD_Stv_ hermaphrodites fed an *E. coli* diet supplemented with EtOH control (-) or with fatty acids (+). N ≥ 24 over 3 independent trials; * *p* < 0.05, Welch’s *t*-test. (**C**) Total, mitotic zone, and transition zone germ cells rows in WT PD_Stv_ animals fed an *E. coli* diet supplemented with EtOH control (-) or OA (+) at different developmental stages. N ≥ 35 over 3 independent trials; no significant differences, one-way ANOVA with Dunnett’s multiple comparison test. (**D**) Representative images of the germline cell rows quantified in (**C**). Yellow and green dotted lines demarcate the mitotic zone and transition zone, respectively. (**E**) Brood sizes of WT PD_Stv_ hermaphrodites fed an *E. coli* diet supplemented with EtOH control (-) or OA (+). N ≥ 45 over five independent trials; ** *p* < 0.01, **** *p* < 0.0001, one-way ANOVA with Dunnett’s multiple comparison test. (**F**) Brood sizes of WT CON hermaphrodites fed an *E. coli* diet supplemented with EtOH control (-) or OA (+). CON animals were transferred at 24h (L1) or 38h (L2) after egg laying. N ≥ 26 over three independent trials; * *p* < 0.05, **** *p* < 0.0001, one-way ANOVA with Dunnett’s multiple comparison test. Graphs indicate mean ± SEM. Additional data provided in Tables S1, S2, and S3.

We showed previously that PD_Stv_ adults have delayed timing for the onset of germline proliferation after dauer exit, which contributes to their lower brood sizes compared to CON adults (Ow et al. 2018). Since *C. elegans* reproduction is sperm-limited, a delay in the onset of germline proliferation leading to spermatogenesis would decrease number of sperm produced, thus decreasing the number of potential progenies resulting from self-fertilization (Byerly, Cassada, and Russell 1976; Ward and Carrel 1979). We next asked whether OA dietary supplementation eliminated the delay in the onset of germline proliferation in PD_Stv_ animals by counting the number of germ cell rows in larvae developmentally synchronized by their vulva development (see Methods). Animals were fed diets that contained either no OA, OA from hatching until dauer, starting at dauer recovery, or throughout life. We found that the OA diet supplementation at any point in life had no significant effect on the number of germ cell rows compared to the control diet (Figure 1C,D). These results indicate that OA-dependent increase in fecundity is acting through a different mechanism than the regulation of the onset of germline proliferation in PD_Stv_ animals.

To determine when OA was required during development to result in an increase in brood size, we counted the brood sizes of PD_Stv_ animals given the same feeding scheme as described above for counting germ cell rows. We found that supplementation after dauer rescue was sufficient to obtain similar brood sizes as supplementation during the entire lifespan. However, dietary supplementation from hatching until the dauer stage did not increase progeny number, suggesting that the amount of lipids stored during dauer does not affect fecundity (Figure 1E). Instead, these results indicate that lipid metabolism after dauer recovery to adulthood influences progeny number. This model is consistent with our previous study showing that NHR-49, which transcriptionally regulates the desaturase *fat-5, -6 and -7* genes (Van Gilst, Hadjivassiliou, and Yamamoto 2005; Gilst et al. 2005), regulates PD_stv_ brood size at a later stage in development without altering the onset of germline proliferation timing. Interestingly, we have shown previously that the OA-dependent increase in PD_Stv_ fecundity is NHR-49 independent, indicating that OA is acting through a different mechanism than that identified to regulate brood size in PD_Stv_ adults (Ow, Nichitean, and Hall 2021).

Given the previous result, we next asked whether the OA-dependent increase in fecundity was limited to PD_stv_ animals. We counted the brood sizes of CON animals fed a control diet, OA from hatching to early larval stage, early larval stage to adulthood, or throughout the lifespan. To emulate the PD_stv_ OA feeding paradigm in continuously growing animals, we transferred the CON larvae at either 24 hours (approximately L1 larval stage) or 38 hours (approximately L2 larval stage) after egg laying. We found that animals fed a diet supplemented with OA at or before the L2 larval stage displayed an increase in brood size similar to animals fed OA throughout life (Figure 1F). Altogether, our results indicate the OA has the special property of positively regulating fecundity regardless of the environmental history of the animal.

### DAF-12 steroid signaling pathway is required for OA dependent increase in fecundity

We have previously shown that somatic aging pathways, including the DAF-12 steroid signaling pathway, were required to regulate PD_stv_ decreased fecundity compared to CON adults. Additionally, we showed that DAF-12 played a role in PD_Stv_ intestinal lipid storage and negatively regulated vitellogenesis (Ow, Nichitean, and Hall 2021). Thus, we tested whether DAF-12 activity was also required for the OA-dependent increase in brood size by feeding strains with mutations in *daf-12* or dafachronic acid biosynthesis genes a control or OA-supplemented diet. We observed that two mutant alleles of *daf-12* that disrupt its dafachronic acid binding activity, *rh284* and *rh285*, eliminated the OA-dependent increase in brood size (Figure 2A). Mutations in the DA biosynthesis genes, *daf-9* and *hsd-1*, also abrogated the increased brood size phenotype, whereas strains with mutations in *daf-36* and *dhs-16* continued to exhibit the OA-dependent increase in progeny similar to wild type (Figure 2A). Notably, DAF- 9, DAF-36, and DHS-16, but not HSD-1, were required for the decreased PD_Stv_ brood size compared to CON adults, providing further support that OA is acting through a separate pathway while still modulating DAF-12 activity (Ow, Nichitean, and Hall 2021).

**Figure 2.**
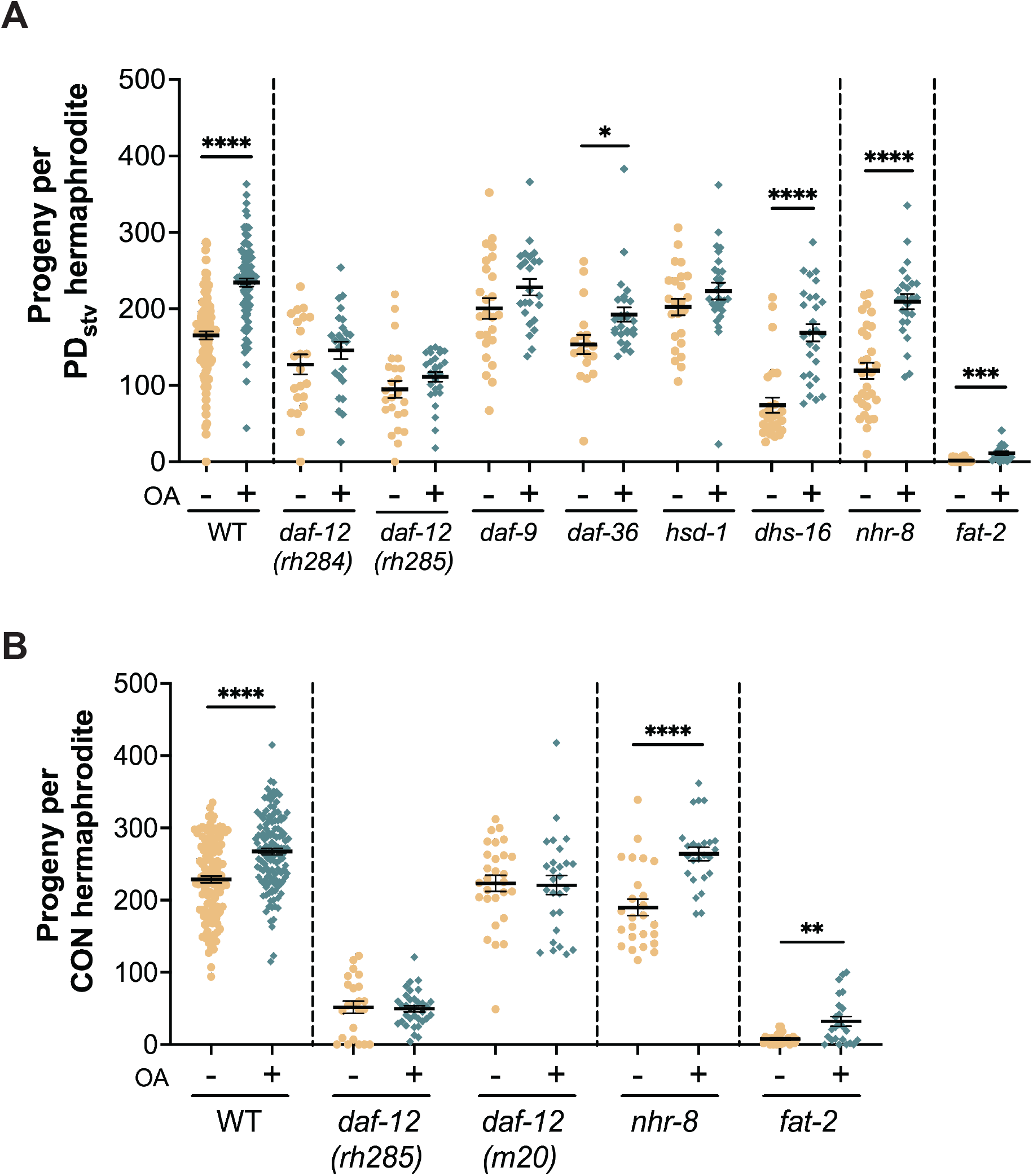
DAF-12 steroid hormone signaling is required for the OA-dependent increase in fecundity. Brood sizes for PD_Stv_ (**A**) and CON (**B**) hermaphrodites fed an *E. coli* diet supplemented with EtOH (-) control or OA (+). Strains include WT, *daf-12(rh284), daf-12(rh285), daf-12(m20), daf-9(rh50), daf-36(k114), hsd-1(mg433), dhs-16(tm1890), nhr-8(ok186)* and *fat-2(wa17)*. N ≥ 18 over three independent trials; * *p* < 0.05, ** *p* < 0.01, *** *p* < 0.001, **** *p* < 0.0001, Welch’s *t* test. Graphs indicate mean ± SEM. Additional data are provided in Figure S1 and Table S4.

Given the role of DAF-12 in regulating OA-dependent brood sizes in PD_Stv_ adults, we also tested whether steroid signaling regulates the same phenotype in CON adults. We observed the brood size of *daf-12(rh285)* mutant CON adults did not exhibit an OA-dependent increase in progeny, indicating that a similar mechanism is regulating the brood size phenotype in both populations (Figure 2B). The *rh284* and *rh285* alleles of *daf-12* are partial loss-of-function mutations that disrupt the ligand binding domain, but still allow for DAF-12 to promote dauer formation in stress conditions. In contrast, the *m20* allele affects the DNA binding domain of the DAF-12 and is considered a null allele. Thus, *daf-12(m20)* animals are dauer deficient (Antebi et al. 2000). To test whether DA driven DAF-12 activity was required for the OA-dependent brood size increase, or whether DAF-12 when bound to its co-repressor DIN-1 specifically inhibits the OA phenotypes, we counted the progeny of *daf-12(m20)* CON adults with and without OA dietary supplementation. We reasoned that if DAF-12 and DIN-1 were acting to suppress the increase in brood size, then the null allele would exhibit a brood size similar to wild type in this experiment. We found that *daf-12(m20)* CON adults do not exhibit the increase in OA-dependent progeny, providing further evidence that DA driven DAF-12 activity is required for this phenotype. The closest homolog of DAF-12 is NHR-8, which is required for cholesterol and bile acid-like DA steroid homeostasis, by regulating *daf-36* and cholesterol availability and transport. Additionally, NHR-8 plays a role in fatty acid metabolism by modulating *fat-5, fat-7, fat-2* and *elo-1* expression, however its effect on fatty acid composition is independent of DAF-12 (Magner et al. 2013). We observed that the OA dependent increase in brood size persisted in *nhr-8(ok186)* CON and PD_Stv_ adults, indicating that NHR-8 is not required for the phenotype (Figure 2). Together, our results indicate that DA driven DAF-12 activity promotes the increased fecundity in the presence of OA regardless of developmental history.

### OA promotes expression of FAT-7 via DAF-12 activity

Lipid metabolism and reproduction are importantly linked in *C. elegans*, such that lipids stored in the soma are transported to the germ line via a process called vitellogenesis (reviewed in (Romano et al. 2004; M. F. Perez and Lehner 2019)). We found previously that the 1′9 desaturase gene, *fat-6,* and Δ12 desaturase, *fat-2,* are required for the observed PD_Stv_ decrease in fecundity compared to CON adults (Ow, Nichitean, and Hall 2021). When we tested whether the 1′9 desaturase genes were also required for the OA-induced increase in progeny, double mutant strains, *fat-5; fat-7* and *fat-6; fat-7*, no longer exhibited the phenotype, suggesting that FAT-7 is required to regulate fecundity in response to OA (Ow, Nichitean, and Hall 2021). We also tested whether FAT-2 was required and found that *fat-2(wa17)* CON and PD_Stv_ animals continued to exhibit a significantly increased brood size with OA supplementation, despite the overall lowered fecundity of the strain (Figure 2). Thus far, *fat-7* is the only lipid metabolism gene we have identified that is required for OA induced phenotypes.

In the fatty acid desaturase pathway, FAT-7 is a stearoyl-CoA desaturase which uses stearic acid as a substrate to produce OA (Figure 1A) (Watts and Browse 2000); thus, our observation that FAT-7 is required to respond to OA supplementation was unexpected. In addition, NHR-49 upregulates the expression of fatty acid biosynthesis genes, including *fat-7*, but we showed that NHR-49 is not required for the OA increase in progeny (Gilst et al. 2005; Ow, Nichitean, and Hall 2021). To further investigate this novel role for FAT-7, we examined whether DAF-12 and FAT-7 may be acting in the same pathway in response to OA by quantifying the expression of a *fat-7p::fat-7::gfp* reporter transgene in CON and PD_Stv_ adults in wild type and *daf-12* strains. In our previous transcriptional profiling experiment, we found that *fat-7* mRNA levels were 5-fold upregulated in PD_Stv_ adults compared to CON adults (Ow et al. 2018). When we quantified GFP expression in the whole intestine of wild type PD_Stv_ and CON adults on a control diet, we found a significantly higher expression in PD_Stv_ intestines, consistent with our RNA-seq data. Interestingly, GFP expression of the transgene significantly increased in both PD_Stv_ and CON adults fed a diet supplemented with OA compared to the control diet (Figure 3). When we repeated the same experiment in *daf-12(rh285)* adults, we again observed an increase in GFP expression in PD_Stv_ intestines compared to CON on the control diet, indicating that DAF-12 is not required for the differential expression of *fat-7* due to developmental history (Figure 3). However, PD_Stv_ and CON intestines no longer exhibited an increase in GFP expression when fed a diet with OA supplementation in the *daf-12* mutants (Figure 3). To confirm our GFP expression quantification, we used qRT-PCR to measure the levels of *fat-7* mRNA present in wildtype and *daf-12(rh285)* PD_Stv_ adults which were given control or OA supplemented diets. Overall, our results matched the trend in *fat-7::gfp* expression that we quantified in the intestine, despite measuring mRNA levels in whole animals (Figure S2). Based on these results, we conclude that *fat-7* expression is increased in animals fed an OA supplemented diet in a DAF-12 dependent manner.

**Figure 3.**
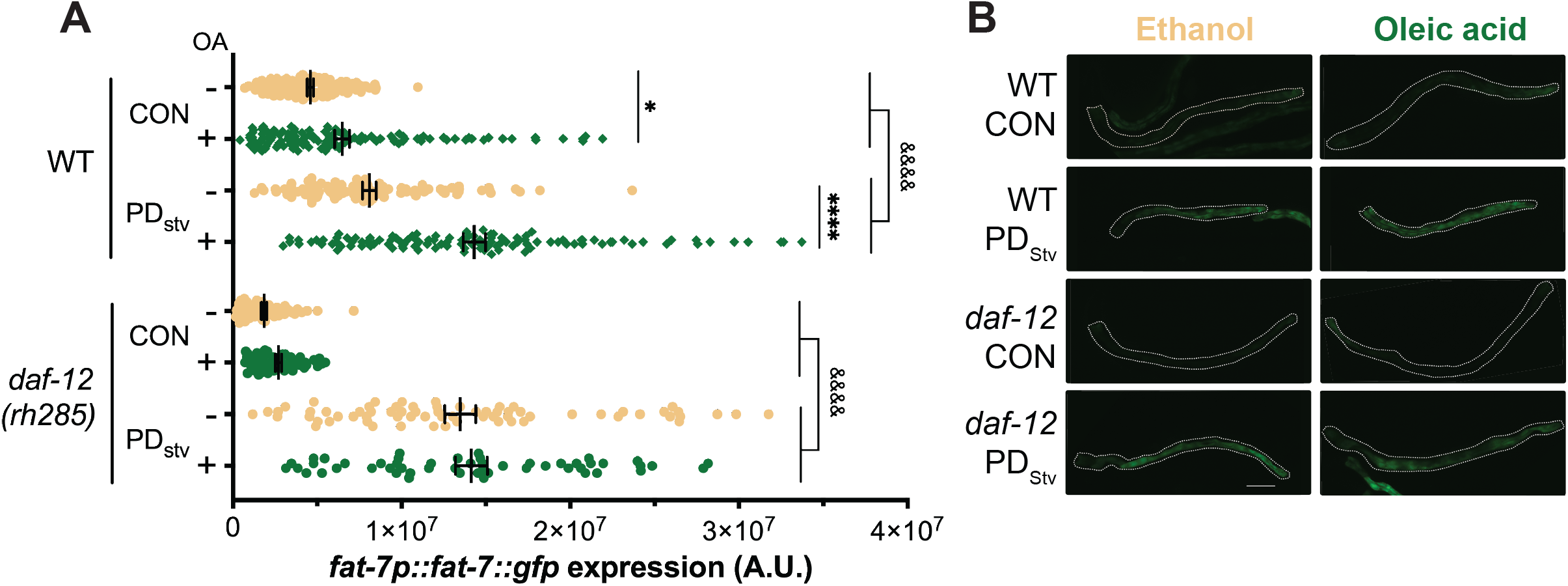
DAF-12 positively regulates *fat-7* expression in response to OA. (**A**) Corrected total cell fluorescence (CTCF) of intestines of *fat-7p::fat-7::gfp* WT and *daf-12(rh285)* CON and PD_Stv_ hermaphrodites fed a diet of *E. coli* supplemented with EtOH control (-) or OA (+). N ≥ 52 over three independent trials; * *p* < 0.05, **** *p* < 0.0001 comparing different diets, ^&&&&^ *p* < 0.0001 comparing CON and PD_Stv_, one-way ANOVA with Šídák’s multiple comparisons test. Graphs indicate mean ± SEM. (**B**) Representative images of animals quantified in (**A**). Dotted line demarcates the intestine. Additional data is provided in Figure S2 and Table S5.

### Vitellogen expression is increased after OA supplementation

Given that we have demonstrated the increased expression of *fat-7* after OA supplementation via DAF-12 (Figure 3), and the negative regulation of vitellogenesis by DAF-12 (Ow, Nichitean, and Hall 2021), we next investigated how OA supplementation may impact vitellogenesis. Vitellogens are yolk proteins expressed in the intestine and taken up by the oocytes through receptor mediated endocytosis (Kimble and Sharrock 1983; Grant and Hirsh 1999; D. H. Hall et al. 1999). First, we examined the expression levels and localization of vitellogen VIT-2. Vitellogens are among the highest expressed protein-coding genes in young adults, and *vit-2*, which encodes a 170 kd polypeptide is the most studied vitellogen of the six genes (Kimble and Sharrock 1983; Nostrand et al. 2013; Goszczynski et al. 2016). Using an integrated *vit-2::gfp* fusion transgene, we quantified GFP expression in the intestine and oocytes of wildtype CON hermaphrodites fed a control or OA-supplemented diet (Figure 4A). In the intestine, GFP was significantly higher in hermaphrodites fed a diet supplemented with OA compared to the control diet, indicating an increase in *vit-2* expression in response to OA (Figure 4B, C). In each of the –3 to –1 oocytes, we also observed a significant increase in GFP levels in OA supplemented hermaphrodites compared those fed a control diet (Figure 4B, C), suggesting that adults fed OA have increased vitellogenesis. Next, we asked whether the increased expression of *vit-2::gfp* after OA supplementation was dependent upon DA driven DAF-12 activity by performing intestinal GFP quantification in animals with RNAi knockdown of *daf-12* and *daf-9*. We were able to replicate the increase in *vit-2::gfp* expression with OA supplementation compared to the control diet in the empty vector RNAi control (Figure 4D). However, the OA-dependent increase in *vit-2* expression was eliminated in the *daf-12* knockdown and was significantly reversed in the *daf-9* knockdown (Figure 4D). While we were not able to recapitulate the increased GFP localization in the –1 and –2 oocytes with OA supplementation using the RNAi protocol, we continued to detect a significant decrease in GFP in animals supplemented with OA with *daf-9* knockdown (Figure S3). These results indicate that DA dependent DAF-12 activity is necessary for increased *vit-2* expression, and possibly localization into oocytes, after dietary OA supplementation.

**Figure 4.**
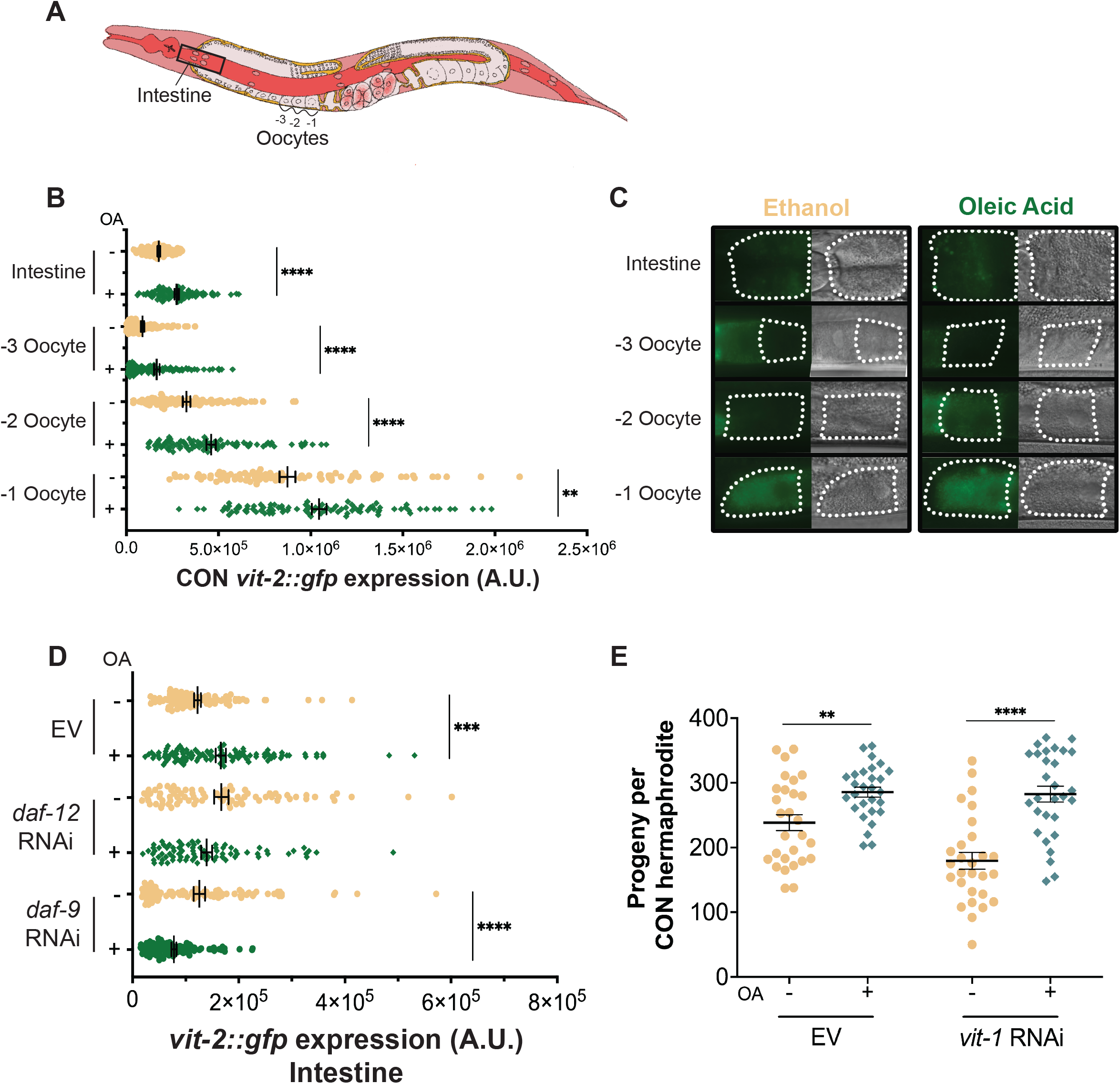
DA dependent DAF-12 activity is required for increased in *vit-2* expression with OA supplementation. (**A**) Diagram of an adult hermaphrodite, indicating the area of the intestine and the oocytes where GFP was quantified. (**B**) CTCF of WT CON hermaphrodites expressing the *vit-2::gfp* transgene fed an *E. coli* diet supplemented with EtOH control (-) or OA (+). N ≥ 87 over three independent trials. (**C**) Representative DIC and fluorescence images of areas quantified in (**B**). (**D**) Intestinal CTCF of WT CON hermaphrodites expressing the *vit-2::gfp* transgene after feeding RNAi with empty vector (EV) control, *f11a1.3* (*daf-12*), and *t13c5.1* (*daf-9*). Animals were fed an *E. coli* diet supplemented with EtOH control (-) or OA (+). N ≥ 78 over three independent trials. (**E**) Brood sizes of WT CON hermaphrodites following feeding RNAi with *k09f5.2 (vit-1)* or empty vector (EV) control. Animals were fed an *E. coli* diet supplemented with EtOH control (-) or OA (+). N ≥ 28 over three independent trials. Graphs indicate mean ± SEM. ** *p* < 0.01, *** *p* < 0.001, **** *p* < 0.0001, Welch’s *t* test. Additional data provided in Figure S4 and Tables S7, S8, and S9.

During our experiments, we noted that the *vit-2::gfp* strain exhibited altered phenotypes relevant to our study, including being dauer deficient and mating deficient. Others have documented additional phenotypes for this strain, including an increase in *vit-2* mRNA compared to wildtype, as well as lower embryo content and brood size (Seah et al. 2016; Erdmann, Abraham, and Hundley, n.d.). Considering that the *vit-2::gfp* transgene may complicate our interpretations of results described above, we tested the hypothesis that vitellogenesis was required for the increased brood size phenotype with OA dietary supplementation using RNAi to knockdown expression of *vit-1* in wildtype hermaphrodites. Knockdown of *vit-1* via RNAi also results in decreased expression of *vit-2/3/4/5* genes due to the high degree of homology within the gene family (Spieth and Blumenthal 1985; Seah et al. 2016; Ow, Nichitean, and Hall 2021). With the empty vector control, we observed the expected increase in brood size with OA supplementation compared to the control diet (Figure 4E). However, knockdown of the *vit* genes failed to eliminate the increased brood size on the OA supplemented diet (Figure 4E). Together, these results suggest that vitellogenesis is increased after OA dietary supplementation, but it is not required for the OA-dependent increase in brood size. The increased expression of *vit-2::gfp* after OA supplementation in hermaphrodites may serve to enhance the survival or adaptability of their progeny to their environment.

### OA rescues germline ferroptosis in PD_Stv_ adults

Thus far, we have demonstrated that dietary supplementation of OA alters metabolism and vitellogenesis in a DAF-12 dependent manner. We next sought to explore the developmental mechanism of how these changes in physiology lead to an increase in fecundity by examining germ cell dynamics. Our results indicate that OA acts after the onset of germline proliferation to regulate the brood size; thus, we hypothesized that OA increases germ cell survival in hermaphrodites. Apoptosis is a well-characterized form of programmed cell death that commonly occurs during the pachytene stage of germ cell development and can be visualized with a variety of molecular tools (Lant and Derry 2014). Approximately half of all germ cells die by stereotypical apoptotic mechanisms in adult hermaphrodites, with one or two cells undergoing apoptosis at any given time point during reproduction (Gumienny et al. 1999; Jaramillo-Lambert et al. 2007). To test whether OA alters apoptosis rates in the germ line, we used two common methods, acridine orange and an *act-5::yfp* transgene, to detect and count cell corpses in CON and PD_Stv_ adult hermaphrodites that were fed a control or OA-supplemented diet. Acridine orange is an intercalating dye that stains DNA of cells undergoing late stages of apoptosis, whereas the *act-5::yfp* transgene is expressed in the gonadal sheath cells and is visualized as “halos” during the engulfment of cell corpses (Lant and Derry 2014). Using both methods, we observed no significant differences in numbers of germline apoptotic cells comparing CON and PD_Stv_ populations or comparing control or OA supplemented diets within a population (Figure 5A, B). We conclude that OA does not increase progeny number by inhibiting apoptosis in the germ line.

**Figure 5.**
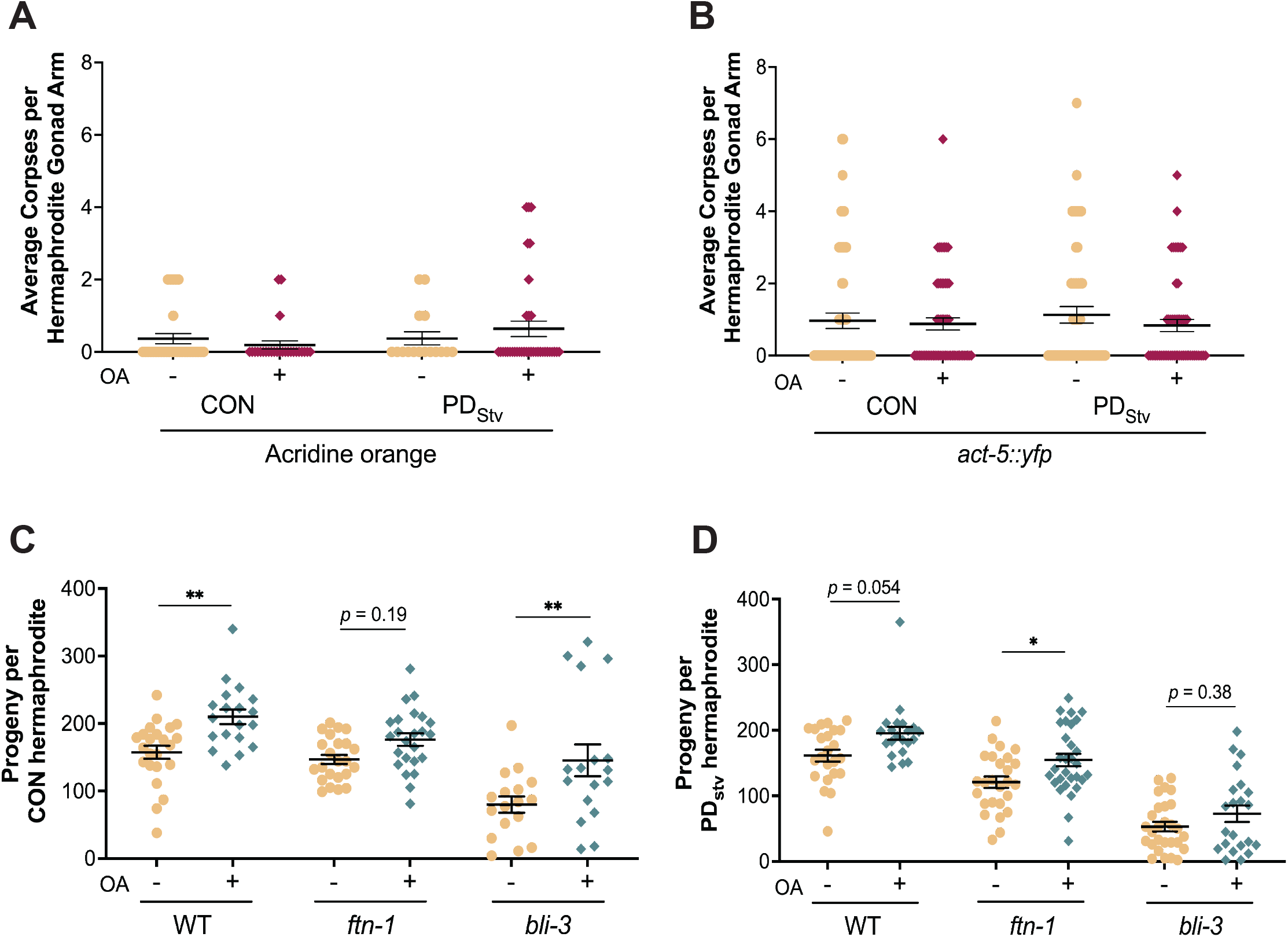
OA rescues germline ferroptosis in PD_Stv_ adults. (**A, B**) Average cell corpses per gonad arm of WT CON and PD_Stv_ hermaphrodites fed an *E. coli* diet supplemented with EtOH control (-) or OA (+) visualized using (**A**) acridine orange dye and (**B**) *act-5::yfp* expression. N ≥ 27 animals per condition over 3 independent trials. No significant differences, one-way ANOVA with Kruskal-Wallis *post hoc* test. (**C, D**) Brood sizes of (**C**) CON and (**D**) PD_Stv_ hermaphrodites fed an *E. coli* diet supplemented with EtOH control (-) or OA (+) for WT, *ftn-1(ok1625)* and *bli-3(e767)* strains. Graphs indicate mean ± SEM. N ≥ 17 animals per condition over 3 independent trials; * *p* < 0.05, ** *p* < 0.01, two-way ANOVA. Additional data provided in Tables S10, S11, and S12.

A second, less characterized form of germ cell death is iron-mediated ferroptosis. Ferroptosis occurs when increased oxidative stress results in incorporation of lipid peroxides in cell membranes, leading to membrane thinning and formation of pores resulting in cell death (Agmon et al 2018). Cell membranes with increased proportions of PUFAs are more susceptible to cell death via ferroptosis: a previous study supplementing the diet of *C. elegans* hermaphrodites with PUFA 20:3n-6 (DGLA) resulted in sterility due to massive cell death in the germ line (Watts and Browse 2006; M. A. Perez et al. 2020). Interestingly, dietary supplementation with both OA and DGLA rescued the sterility phenotype of DGLA alone, indicating that OA was able to block ferroptosis of germ cells (M. A. Perez et al. 2020). We showed previously using lipid profiling that PD_Stv_ adults had a significantly increased level of DGLA compared to CON adults (Ow, Nichitean, and Hall 2021). Thus, we tested the hypothesis that PD_Stv_ adults have increased ferroptosis in their germlines contributing to their decreased fecundity, which is rescued by OA dietary supplementation. To test our hypothesis, we examined the brood sizes of strains carrying mutations in genes with known functions that positively or negatively regulate the incidence of ferroptosis that have been fed control or OA supplemented diets. FTN-1 is the *C. elegans* homolog of ferratin, an iron sequestering protein, and mutations in *ftn-1* should increase ferroptosis due to an excess of ferrous iron generating reactive oxygen species (ROS) and promoting lipid peroxidation (Gourley et al. 2003; Kim et al. 2004; Valentini et al. 2012; M. A. Perez et al. 2020). In contrast, BLI-3 is an NADPH oxidase and the closest *C. elegans* homolog to mammalian NOX, and mutations in *bli-3* are pleiotropic due to its functions in cuticle development and the production of ROS during an immune response (Edens et al. 2001; Simmer et al. 2003; Chávez, Mohri-Shiomi, and Garsin 2009; Hoeven et al. 2015). Additionally, *bli-3* mutants display resistance to DGLA induced sterility, thus making it a positive regulator of ferroptosis (M. A. Perez et al. 2020). We predicted that mutations in *ftn-1* and *bli-3* would increase or decrease the levels of ferroptosis, respectively, and that OA would have a more significant impact on the brood size of *ftn-1* animals compared to *bli-3*. A two-way ANOVA of our CON and PD_Stv_ brood sizes for wild-type, *ftn-1*, and *bli-3* strains indicated that genotype and diet were significant sources of variation, but there was no interaction between the two. For CON adults, we observed a significant increase in wild type progeny number with OA supplementation compared to the control diet as expected (Figure 5C). However, *ftn-1* and *bli-3* CON adults exhibited the opposite phenotypes from our prediction, with OA supplementation having no effect on *ftn-1* progeny number but significantly increasing the brood size of *bli-3* adults (Figure 5C). These results suggest that ferroptosis may be less physiologically relevant in CON adult germ lines, and that OA may be acting through a different mechanism to increase brood size in this population. In contrast, the PD_Stv_ adults exhibited brood sizes as we predicted if ferroptosis were occurring in the germ line. Wild type animals exhibited a trend of greater brood sizes with the OA supplemented diet compared to controls, although not quite significant in these trials. *Ftn-1* mutants have significantly increased brood sizes with OA supplemented diets, whereas OA does not affect *bli-3* progeny number (Figure 5D). These results suggest that PD_Stv_ and CON adult germ lines have different levels of intrinsic ferroptosis and that OA may act through different mechanisms to increase brood size in these populations. Further experiments with more directed tools to study ferroptosis will be necessary to further explore these differences in CON and PD_Stv_ adults.

### OA decouples reproduction and longevity in adults

In *C. elegans,* longevity and reproduction are thought to have an inverse proportional relationship. Larvae that have no germline progenitor cells or lack functional GLP-1/Notch protein have extended lifespans, which are dependent on the DAF-12 and fatty acid metabolism pathways (Hsin and Kenyon 1999; Arantes-Oliveira et al 2002; Yamawaki et al. 2010; McCormick et al. 2012; Mahanti et al. 2014). These results have led to a model whereby the absence of germ cells increases the resources for somatic maintenance, allowing for an extended lifespan. OA has also been demonstrated to play an important role in lifespan regulation. Mutations in *fat-6* and *fat-7* abrogate the increased longevity of *glp-1* mutants, which can be restored with dietary OA supplementation (Goudeau et al. 2011). Additionally, hermaphrodites that have mated with males have a significantly reduced lifespan compared to unmated controls, which can be rescued through dietary OA supplementation (Choi et al. 2021). However, conflicting results on longevity have been observed when OA is added directly to the plate agar (Goudeau et al. 2011; Ratnappan et al. 2014; Lee et al. 2015; Han et al. 2017; L. Zhou et al. 2021).

Given the ability of OA to extend lifespan in certain circumstances, and our results demonstrating that OA can increase progeny number, we sought to investigate the consequences on OA on longevity using our experimental system. To this end, we examined the adult lifespan of wildtype and *daf-12(rh285)* CON and PD_Stv_ adults given a control diet, OA supplementation in early development, OA supplementation in late development, or OA supplementation throughout their lifespan. First, we observed that the lifespans of CON and PD_stv_ adults were affected differently depending on the timing of OA supplementation during development. For CON adults, lifespan was significantly increased when OA supplementation occurred either before or after L2 stage, which suggests that intermittent OA supplementation helps mitigate the stress of reproduction on somatic maintenance (Figure 6A). In addition, the increased lifespan of CON adults is independent of their increased fecundity, since we have shown that only animals that have OA supplementation after L2 stage have increased brood size (Figure 1F). CON adults fed dietary OA throughout life exhibited longevity similar to the control diet, consistent with previous findings (Goudeau et al. 2011; Ratnappan et al. 2014; Lee et al. 2015; Choi et al. 2021). *Daf-12(rh285)* CON adults no longer exhibited the increased longevity with intermittent OA, but did show a slight, but significant, increase with continued OA compared to the control diet (Figure 6B), indicating that DAF-12 plays a role in the OA- depended increase in lifespan of CON animals.

**Figure 6.**
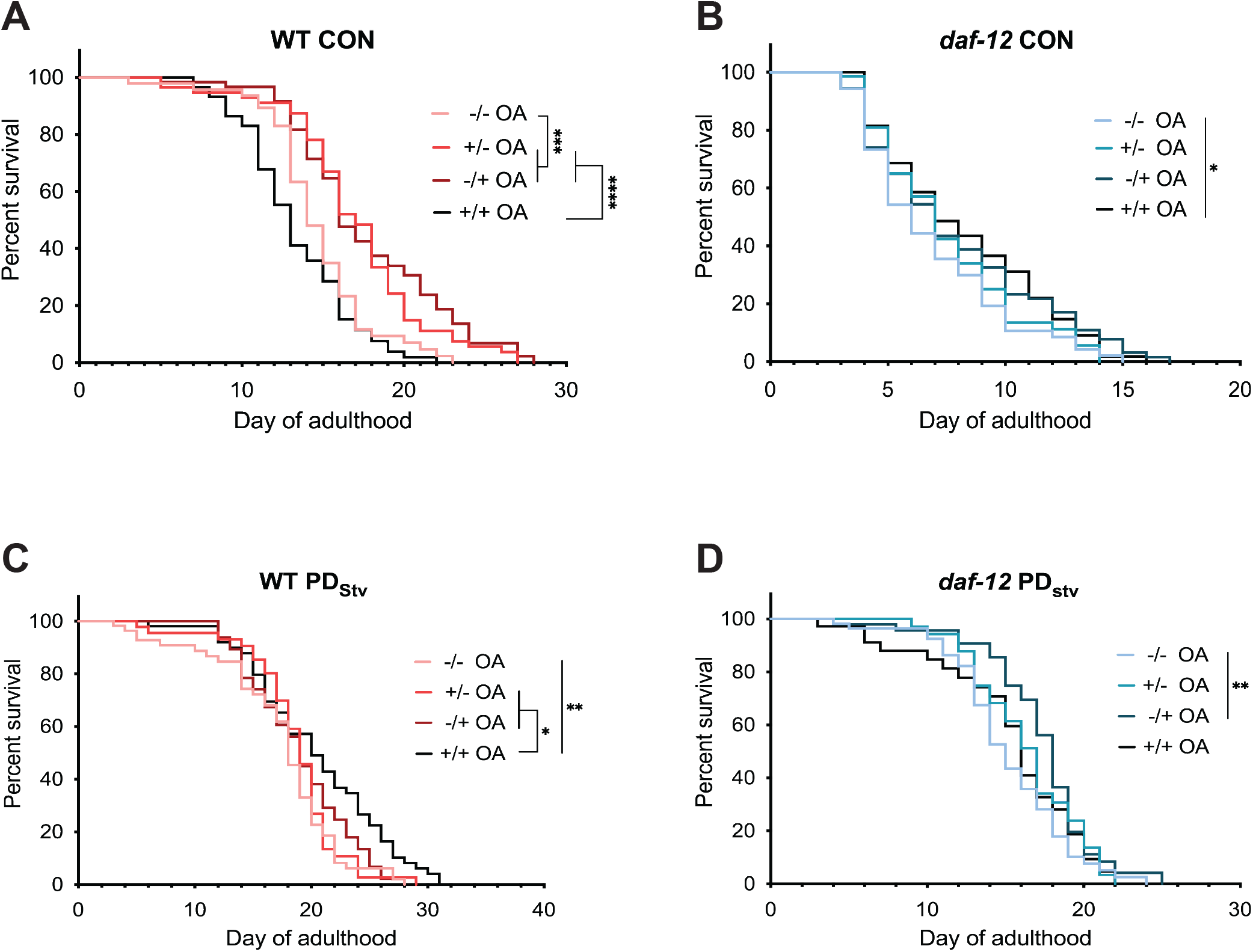
OA increases longevity in *C. elegans* adults. Adult lifespan assays of (**A**) WT CON, (**B**) *daf-12(rh285)* CON, (**C**) WT PD_Stv_ and (**D**) *daf-12(rh285)* PD_Stv_ hermaphrodites. Animals were fed an *E. coli* diet supplemented only with EtOH control (-/-), OA only from hatching until L2 or dauer (+/-), OA only from L2 or dauer until adulthood (-/+), or OA throughout lifespan (+/+). N ≥ 26 over 3 independent trials; * *p* < 0.05, ** *p* < 0.01, *** *p* < 0.001, **** *p* < 0.0001, log-rank (Mantel-Cox) test. Additional data provided in Table S13.

In contrast, PD_stv_ adults only exhibited significantly increased longevity when fed OA supplementation throughout life (Figure 6C). In the *daf-12* mutants, passage through dauer rescued the reduced longevity of the *daf-12* CON adults (Figure 6B,D). Furthermore, *daf-12* PD_Stv_ adults that were given OA after dauer rescue had a significant increase in the median life span compared to animals fed a control diet (Figure 6D). Together, these results indicate that the timing of OA dietary supplementation timing during larval development can modulate lifespan in a manner that is specific to the animals’ life history. In addition, the lack of correlation between the conditions that extend lifespan and increase fecundity suggests that the mechanisms controlling these life history traits are decoupled and modulated differently by OA supplementation.

## DISCUSSION

In this study, we demonstrate that dietary OA supplementation increases fecundity and lifespan of *C. elegans* adults regardless of the animals’ developmental history. We showed that OA supplementation later in larval development after the onset of germline proliferation (after dauer or L2) was sufficient to manifest the increased fecundity, but the timing of OA supplementation for increased longevity was dependent upon life history of the animals. Both of these OA-induced phenotypes required DAF-12. We showed that dafachronic acid biosynthetic enzymes, DAF-9 and HSD-1, were required for the increased fecundity, in addition to the FAT-7 1′9 desaturase that we found previously (Ow, Nichitean, and Hall 2021). Moreover, we found that dietary supplementation of OA altered the activity of DAF-12 such that it promoted increased expression of *fat-7* and *vit-2*. To understand the developmental mechanism of how OA increases fecundity, we examined cell death in CON and PD_Stv_ adult germ lines. Our results suggest that OA rescues cell death by ferroptosis in PD_Stv_ germlines, but CON fecundity may be increased by a different mechanism. We propose that OA is capable of modulating fecundity and longevity independently of each other while still sharing the common mechanism of altering DAF-12 steroid hormone signaling.

### DAF-12 is the master regulator of OA-dependent phenotypes

One of the common factors between the dauer-dependent brood size decrease and the OA-dependent brood size increase in PD_Stv_ adults is DAF-12. These two phenotypes are at odds with each other, yet by supplementing the diet of PD_Stv_ animals with OA, DAF-12 ultimately rescues its own phenotype. DAF-12 is required for the decreased fecundity of PD_Stv_ adults, partially by delaying the onset of germline proliferation in postdauer larvae (Ow, Nichitean, and Hall 2021). While this phenotype remains unaltered in PD_Stv_ larvae fed dietary OA (Figure 1C), DAF-12 also acts to increase the progeny number in these same animals (Figure 2A). These results introduce the intriguing question of how DAF-12’s activity is altered in the presence of OA. Using alleles of *daf-12* which have a mutation in the ligand binding domain, we were able to determine that the ability of DAF-12 to bind dafachronic acid is key to regulating OA-dependent fecundity (Figure 2B). DAF-12’s activity as a transcription factor is modulated by which DA molecule it is bound to and the life stage of the animal (Mahanti et al. 2014). For example, Δ^4^- DA dependent DAF-12 activity can have distinct effects on longevity depending on environmental factors and tissue specificity. In germline-less adults, DAF-12 and Δ^4^-DA is required for increased longevity. However, during favorable conditions, liganded DAF-12 promotes reproductive growth and a shorter life span (Gerisch et al. 2007). Recent studies have identified numerous new metabolites that differ between *C. elegans* strains and by sex; thus, the possibility exists that OA triggers the biosynthesis of novel DAF-12 ligands that alter its activity accordingly (Fox et al. 2022; Burkhardt et al. 2023).

Given that NHRs commonly function to regulate transcription as heterodimers, another possibility is that DAF-12 activity is modulated by binding with a distinct protein partner that binds to OA as a ligand. In organ culture, oleic acid and linoleic acid were found to be activators of peroxisome proliferator-activated receptor-alpha (PPAR-α) (Hanley et al. 1997). PPAR-α transcription factors are orphaned nuclear receptors that bind to a peroxisome proliferator response element in the promoter region of target genes. PPAR-αs are most highly expressed in tissues with high rates of mitochondrial and peroxisomal fatty acid β-oxidation rates, and they can function as fatty acid sensors and regulate fatty acid metabolism by activating lipid metabolism genes (van Raalte et al. 2004). In mice, SCD1, the homolog of FAT-7, is transcriptionally regulated by PPAR (Miller and Ntambi 1996). The *C. elegans* genome encodes a large gene family of 284 nuclear receptors, the majority of which have unknown ligands (Taubert, Ward, and Yamamoto 2011). Future work will be necessary to determine if either of these possibilities are relevant to how DAF-12 has altered activity after dietary OA supplementation.

This altered activity of DAF-12 in the presence of OA affects the downstream expression of other genes, including *fat-7* and *vit-2* (Figures 3 and 4). In our study, we saw that that FAT-7 was required for the OA-dependent increase in brood size, and that *fat-7* expression is significantly increased with OA supplementation in a DAF-12 dependent manner (Figure 3). The regulation of *fat-7* by DAF-12 appears to be specific to the presence of OA, as the increase in *fat-7* expression levels in PD_Stv_ adults compared to CON adults in the absence of OA is not DAF-12 dependent. Additionally, these results raise the question of why *fat-7* is induced in the presence of OA. FAT-7 is a Δ9 desaturase that modifies stearic acid to form OA (Watts and Browse 2000), so it is unclear why the enzyme required to synthesize OA is upregulated in animals fed OA. Interestingly, other phenotypes rescued by OA, such as mating-induced death in hermaphrodites, also require FAT-7, indicating that this observation is not unique to our study (Han et al. 2017; Choi et al. 2021; L. Zhou et al. 2021). At high temperatures, FAT-7 is important for membrane lipid fluidity during heat adaptation and survival. *Fat-7* expression is decreased at high temperatures, but RNAi knockdown of *fat-7* at 25° C increases its mRNA levels compared to controls, suggesting that FAT-7 expression may be regulated by a complex feedback mechanism (Ma et al. 2015; L. Zhou et al. 2021). Furthermore, FAT-7 is annotated as having a predicted iron ion binding function, which may link the DAF-12, FAT-7, and ferroptosis aspects of our study (Gaudet et al. 2011).

### OA and the connection between fecundity, vitellogenesis, and longevity

The long-standing theory of aging is that the “cost” of germline production deprives the soma of resources, thereby decreasing longevity (Kirkwood et al. 1997; Heininger 2002). Examples of this soma-germline conflict can be found in *Drosophila melanogaster*, *Seychelles warblers* and mice (Smith 1958; Stearns and Kaiser 1996; Cargill et al. 2003; Rogina et al. 2007; Hammers et al. 2013). In *C. elegans*, DAF-12 steroid hormone signaling is key to the evidence supporting the so-called “disposable soma” hypothesis. DA-dependent DAF-12 activity results in adults that experience reproductive development and a shorter lifespan, but the same pathway promotes longevity in the absence of a germ line (Gerisch et al. 2007). Our results in this study indicate that OA can alter DAF-12 activity so that reproduction and lifespan are decoupled. The discrepancy of our data with the somatic aging theory is illustrated by the regulation of vitellogenesis and lifespan. *C. elegans* lifespan is indirectly proportional to vitellogen production, such that decreased expression of *vit-1/-2/-3/-4/-5* genes is associated with increased lifespan, and vice versa (Seah et al. 2016; Sornda et al. 2019; Ow, Nichitean, and Hall 2021). We showed previously that DAF-12 negatively regulates vitellogenesis in CON and PD_Stv_ adults. PD_Stv_ adults have a decreased brood size and increased longevity, despite also having increased vitellogenesis compared to CON adults. When *vit* genes were knocked down in PD_Stv_ adults by RNAi, their lifespan increased even further (Ow, Nichitean, and Hall 2021). In this study, we show that DA-dependent DAF-12 activity increases *vit-2* expression and fecundity after OA dietary supplementation (Figures 1 and 4), both of which have been associated with decreased longevity. However, our results demonstrate that OA dietary supplementation also allows for increased longevity through the activity of DAF-12, despite the augmentation of other “costly” life history traits (Figure 6). All together, these results suggest a model where 1) these pathways (reproduction, longevity, and vitellogenesis) are regulated separately, and their phenotypes are largely correlative in mutant backgrounds or certain environmental conditions, or 2) these pathways are co-regulated and OA alters DAF-12 activity in such a way that it “overrides” the typical programming. Our observation that DAF-12 functions to both decrease and increase brood size in PD_Stv_ adults fed dietary OA supports the second model. Future work will be necessary distinguish between these two models.

In humans, the tradeoff between somatic growth and fecundity is less clear due to factors such as social-economic status, location of residence, and medical history playing significant roles in human physiology. Evidence that OA also improves life history traits in humans is evident from the documented benefits of the Mediterranean diet, where OA is a prominent ingredient found in olive oil. A large study examining the effects of a Mediterranean diet supplemented with olive oil or nuts showed improvements in blood pressure, insulin sensitivity, lipid profiles, lipoprotein particles, inflammation, oxidative stress, and carotid atherosclerosis (Martínez-González et al. 2015). Additional studies more specifically examining the effects of OA dietary supplementation largely boasted decreased inflammation, increased insulin resistance, increased physical activity, and improved mood (Palomer et al. 2018; Kien, Bunn, Tompkins, et al. 2013; Kien, Bunn, Poynter, et al. 2013). These studies support a role for OA in improving the health span of humans. While the effects of dietary OA on human fertility are not well studied, studies in other mammals suggest that OA dietary supplementation may improve obesity-related infertility, but at high levels may have adverse effects on female fertility (e.g. (Yousif et al. 2020; X. Zhou et al. 2022)). Whether the effects we have reported here in *C. elegans* translate to people that are experiencing the effects of metabolic programing, or if they may indicate benefits for all populations, still remain to be determined.

## MATERIALS AND METHODS

### *C. elegans* strains, maintenance, and fatty acid supplementation

The following strains have been used in this study: N2 Bristol (WT); AA82 *daf-12(rh284)* X; AA85 *daf-12(rh285)* X; RG1228 *daf-9(rh50)* X; AA292 *daf-36(k114)* V; GR2063 *hsd-1(mg433)* I; AA1052 *dhs-16(tm1890)* V; AE501 *nhr-8(ok186)* IV; BX26 *fat-2(wa17)* IV; DMS303 *nIs590*[*fat-*7p::fat-7::gfp; lin15(+)] V; SEH438 daf-12(rh285); nIs590[fat-7p::fat-7::gfp; lin15(+)] V; RT130 pwIs23[vit-2::gfp]; WS2170 opIs110[lim-7p::yfp::actin + unc-119(+)] IV; CB767 bli-3(e767) I; RB2603 ftn-1(ok3625) V. The C. elegans strains were maintained at 20°C on NGM plates and fed E. coli OP50 using standard methods when not used for assays (Brenner 1974; Stiernagle 2006).

When the animals were used in assays, they were maintained on peptone-free NGM plates at 20°C. The peptone free plates were seeded with 10X *E. coli* grown in superbroth, which was supplemented with either the fatty acid or an equal concentration of ethanol, and left to grow overnight (Ow, Nichitean, and Hall 2021). Fatty acid supplemented food was made fresh for each assay and was seeded on plates no more than 24 hours before use. The fatty acids used in diet supplementation include: oleic acid (Nu-Chek-Prep, INC.), palmitoleic acid (Sigma), linoleic acid (Sigma), arachidonic acid (Sigma); and eicosapentaenoic acid (Supelco) (O’Rourke et al. 2013; Devkota et al. 2017; Anderson et al. 2019; Ow, Nichitean, and Hall 2021). To obtain CON animals, L4 stage larvae were maintained on NGM plates for two days until adulthood, then placed on peptone-free plates with or without the fatty acid supplementation to lay eggs for 1-2h, after which the adults were removed. The larvae were then either left to grow on the food source until adulthood or they were transferred to the alternative food source at 24h after egg laying (approximately L1) or 38h after egg laying (approximatively L2). To collect starvation-induced dauers, hermaphrodites were maintained on the peptone free plates with the limited food until dauers were observed. After 48 hours, the animals were washed with SDS using standard methods (Karp 2018) and rescued on a fresh peptide-free plate with ethanol control or fatty acid supplemented food.

### Brood size assays

Brood size assays were performed as previously described with some modifications (S. E. Hall et al. 2010). Individual L4 larvae were placed on plates seeded with the desired food type and transferred every two days to fresh peptone free plates with the same food type for a total of 6 days. Hatched progeny were counted for each hermaphrodite to determine total brood size. Results from adults that died, exploded, or dried on the side of the plate during the assay were censored from the final data. Depending on the experimental setup, a *t*-test or ANOVA was used to test for statistical significance using GraphPad Prism (version 9.5).

### Imaging and quantification of GFP expression

CON and PD_Stv_ day one adults were placed on 1% agarose gel pads and anesthetized using sodium azide. Animals were imaged using a Leica DM5500 B microscope with a Hamamatsu camera controller C10600 ORCA-R2. The images were analyzed using ImageJ and the cell total corrected fluorescence (CTCF) was calculated using the formula CTCF = Integrated density of ROI – Integrated density of background selection, when the area of the ROI was identical to the background. Alternatively, when the area of the ROI and background was different, we calculated CTCF as Integrated density of ROI – (Area of ROI *background mean grey value). The CTCF values were inputed into GraphPad Prism (version 9.5) for statistical analysis.

### Feeding RNAi experiments

A 10X concentrated bacterial culture containing the *f11a1.3* (*daf-12*), *t13c5.1* (*daf-9*), *k09f5.2* (*vit-1*), or empty vector RNAi clone from Ahringer library (Kamath et al. 2000) was used to seed NGM plates containing 1 mM IPTG and 50 ug/ml carbenicillin. Adult hermaphrodites were placed on RNAi plates to lay 12-20 eggs, after which the animals were removed. When the F1 generation became reproductive adults, approximately 20 eggs and larvae from the F2 generation were transferred to a new RNAi plate. This step was repeated to obtain the F4 generation. The adults of F4 generation were placed on peptone free plates seeded with *E. coli* supplemented with either ethanol or OA to lay eggs. When the F5 generation reached the L4 larval stage, the animals were used for imaging or brood size assays. As a control, we performed feeding RNAi with the *k09f5.2* (*vit-1*) clone on the animals expressing the *vit-2::gfp* transgene to confirm that the *gfp* was silenced.

### DAPI staining

Counting germ line cell rows was performed as described previously (Ow et al. 2018). Briefly, animals were synchonized based on their vulval morphology using Nomarski DIC microscopy at 630X, followed by DAPI staining using the standard protocol. The images were taken using a Leica DM5500B microscope with a Hamamatsu camera controller C10600 ORCA-R2. Cell row counts were performed using ImageJ. The boundary between mitotic and transition zones was identified where at least two cells in a row showed the crescent-shaped nuclear morphology (Shakes et al. 2009).

### Acridine Orange staining

Thirty to forty L4 larvae of each condition (CON and PD_Stv_) were maintained on peptone-free plates with food supplemented with either OA or ethanol. After 24 hours, 75 μg/mL acridine orange was applied to just cover the bacterial lawn, and the plates were incubated in the dark for 1 hr with the lids ajar. Animals were then moved to fresh control or OA supplemented plates and allowed to incubate in the dark for 3 additional hours. Animals were mounted onto 2% agarose pads in 2-4 μL of 1M sodium azide to score for cell death using the Leica DM5500B microscope at 630X magnification on the GFP filter. Both gonads arms were scored for each animal if possible.

### ACT-5::YFP scoring

Fifty L4 larvae for each condition (CON and PD_Stv_) were picked to plates with either EtOH control or OA supplemented diet and allowed to age 24 hrs. Day 1 adults were then mounted in 4 μL of 1 M of sodium azide on freshly made 2% agarose pads. Animals were scored for apoptotic corpses using the Leica DM5500B microscope at 630X magnification on the YFP filter. Both gonad arms were scored for each animal if possible.

### qRT-PCR

Total RNA from WT and *daf-12(rh285)* 1 day-old adults from four biological replicates cultivated on EtOH control or OA supplemented plates was extracted using Trizol reagent (Sigma) and reverse transcribed using Superscript IV following the manufacturer’s protocol (Invitrogen). Quantitative PCR was performed using SYBR Green Master Mix (Biorad). The experimental mRNA levels were normalized using two somatic genes, *f28b4.3* and *y45f10d.4* (Ow et al. 2018). Relative fold changes of mRNA levels were calculated using the ΔΔCt method. Student’s *t*-test was used to determine statistical significance.

### Lifespan assay

Ten to twenty L4 larval stage animals were placed on individual peptone free plates seeded with *E. coli* OP50 supplemented with EtOH control or OA. Throughout the reproductive period, the animals were moved to new plates every two days. Animal survival was assessed every day by examining for pharynx pumping and movement after gentle touch. Animals that crawled up the side of the plate wall or died by bursting or bagging were censored from the experiment. Statistical analysis was performed using GraphPad Prism version 9.5.

## Supporting information

Supplemental tables with raw data

## ACKOWLEDGMENTS

We are grateful to Eleanor Maine for thoughtful comments on this manuscript. We thank the CGC, which is funded by the NIH Office of Research Infrastructure Programs (P40 OD010440), for providing strains. This work was partially supported by a NIHR01GM129135 grant to S.E.H.

## AUTHOR CONTRIBUTIONS

A.M.N., F.V.C., and S.E.H. conceived and designed experiments. A.M.N. and F.V.C. performed experiments. A.M.N., F.V.C., and S.E.H. analyzed the data. A.M.N., F.V.C., and S.E.H. wrote the manuscript.

## SUPPLEMENTAL MATERIALS LIST

Figure S1 – related to Figure 2

Figure S2 – related to Figure 3

Figure S3 – related to Figure 4

Table S1 – data for Figure 1B

Table S2 – data for Figure 1C

Table S3 – data for Figure 1E and F

Table S4 – data for Figures 2 and S1

Table S5 – data for Figure 3

Table S6 – data for Figure S2

Table S7 – data for Figure 4B

Table S8 – data for Figure 4D

Table S9 – data for Figure 4E

Table S10 – data for Figure 5A

Table S11 – data for Figure 5B

Table S12 – data for Figure 5C and D

Table S13 – data for Figure 6

**Figure S1.**
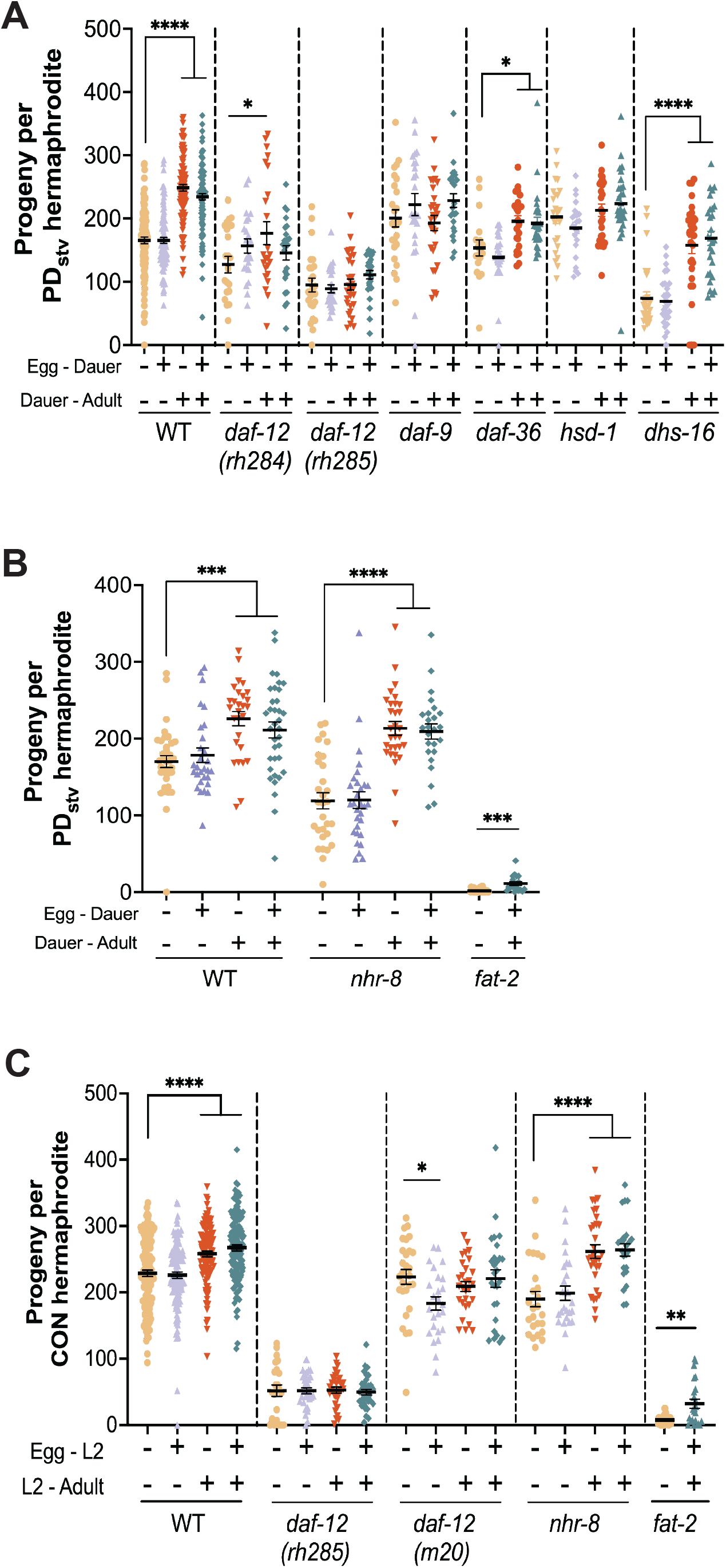
OA dependent increase in fecundity after dauer rescue requires DAF-12. (**A**, **B**) Brood sizes of PD_Stv_ hermaphrodites fed with an *E. coli* diet supplemented with EtOH control (-) or OA (+) for (**A**) WT, *daf-12(rh284)*, *daf-12(rh285)*, *daf-9(rh50), daf-36(k114), hsd-1(mg433), dhs-16(tm1890)* and (**B**) WT, *nhr-8(ok186), fat-2(wa17)* strains. (**C**) Brood sizes of CON hermaphrodites fed a diet as described in (**A,B**) for WT, *daf-12(rh285)*, *daf-12(m20)*, *nhr-8(ok186)*, *fat-2(wa17)* strains. Graphs indicate mean ± SEM. N ≥ 18 over three independent trials; * *p* < 0.05, ** *p* < 0.01, *** *p* < 0.001, **** *p* < 0.0001, one-way ANOVA with Dunnett’s multiple comparison test. Additional data is provided in Table S4.

**Figure S2.**
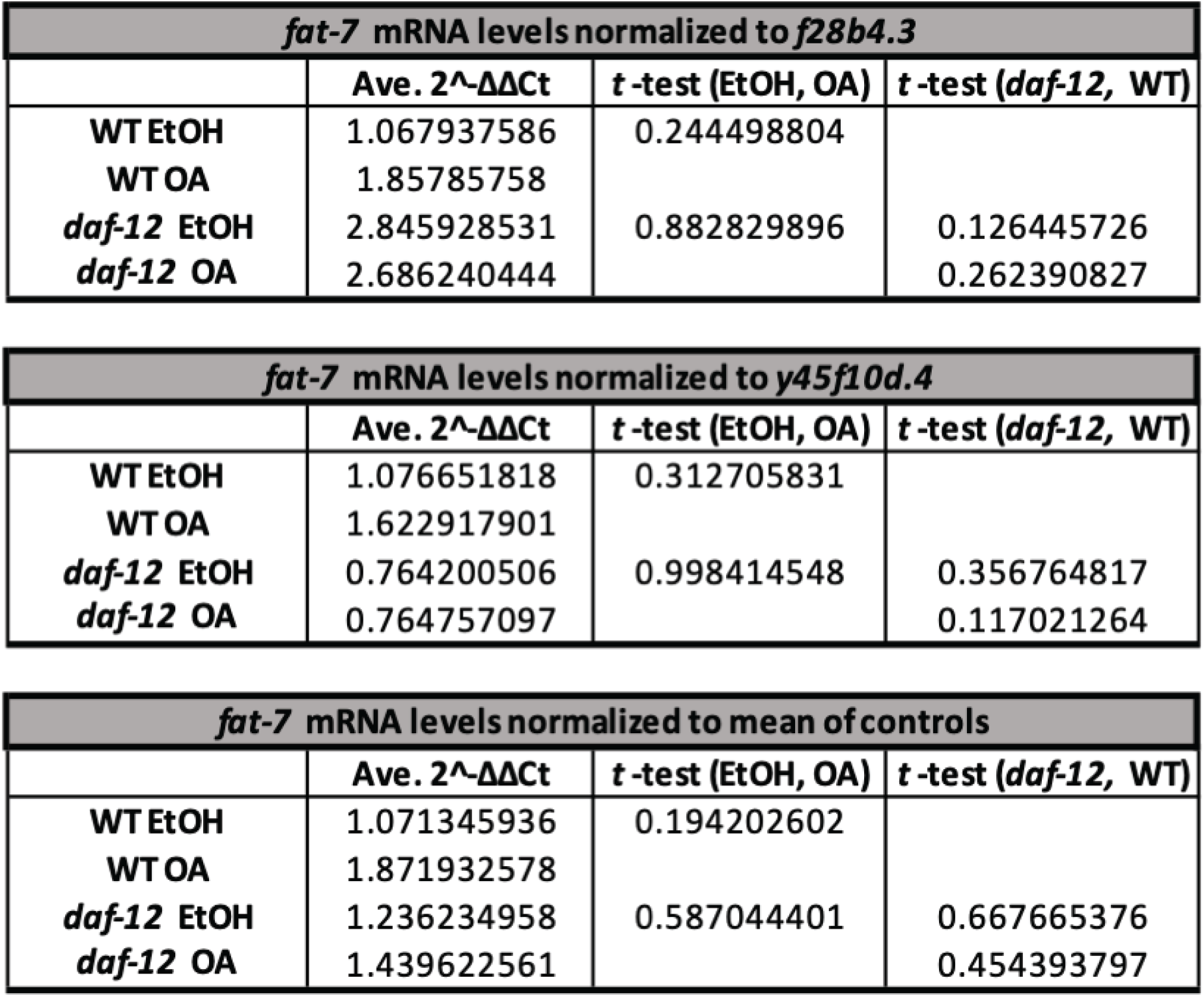
*fat-7* expression increases in whole animals in response to OA. mRNA levels of *fat-7* measured using qRT-PCR analysis by the ΔΔCt method in WT and *daf-12(rh285)* one-day-old adults fed an *E. coli* diet supplemented with EtOH control or OA. 2^^- ΔΔCt^ was normalized to *f28b4.3* and *y45f10d.4* somatic housekeeping genes Ct. Additional data provided in Table S6.

**Figure S3.**
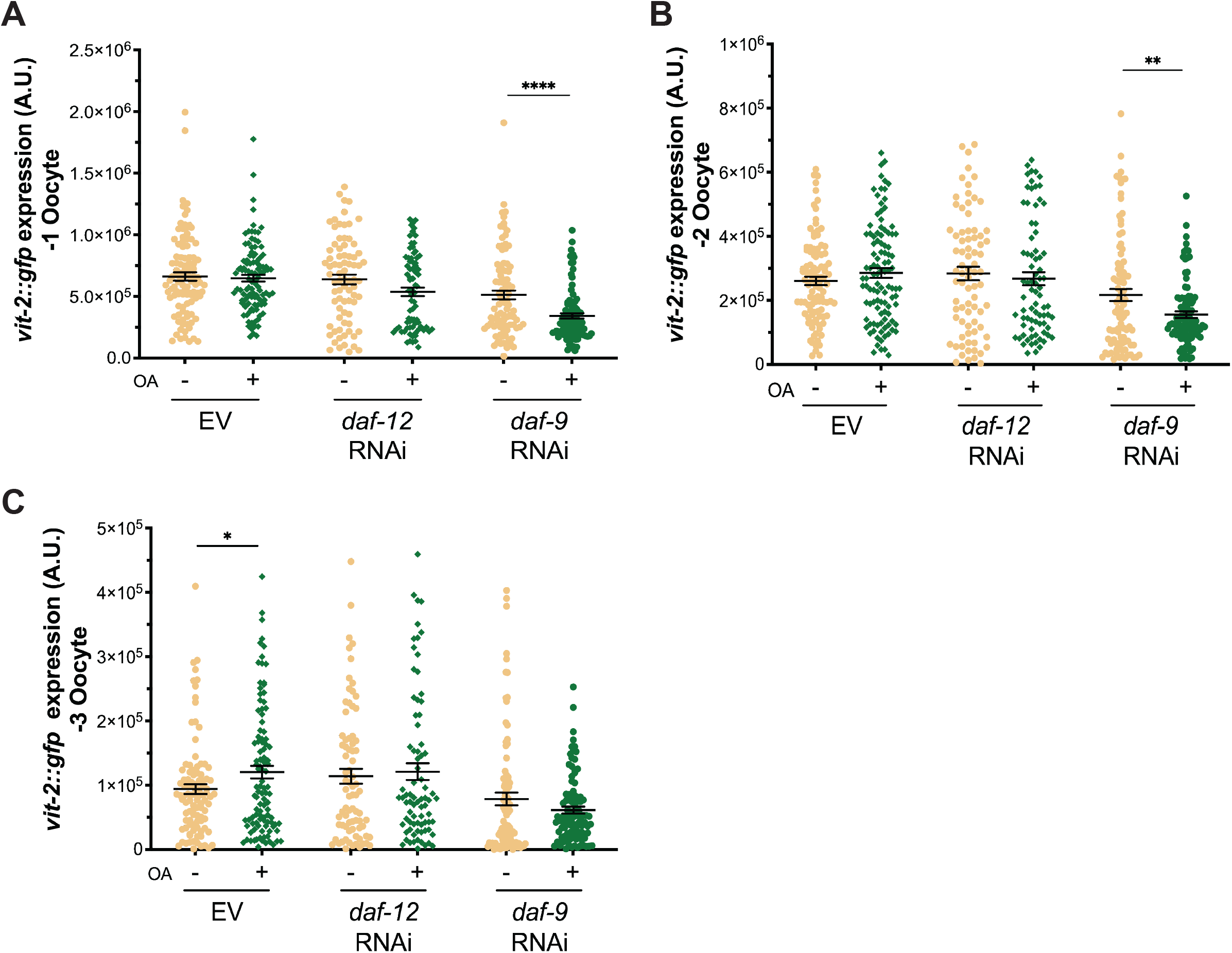
DAF-12 and DAF-9 are required for the increase in *vit-2* expression in response to OA. CTCF of (A) -1 oocytes, (B) -2 oocytes, and (C) -3 oocytes in WT CON hermaphrodites expressing the *vit-2::gfp* transgene following feeding RNAi with *f11a1.3 (daf-12), t13c5.1 (daf-9),* or empty vector (EV) control. Animals were fed an *E. coli* diet supplemented with EtOH control (-) or OA (+). Graphs indicate mean ± SEM. N ≥ 78 over three independent trials; * *p* < 0.05, ** *p* < 0.01, **** *p* < 0.0001, Welch’s *t* test. Additional data provided in Table S8.

